# Morpho-anatomical variation and their phylogenetic implications in native and exotic species of *Pinus* growing in the Indian Himalayas

**DOI:** 10.1101/754804

**Authors:** Lav Singh, Pooja Dixit, Ravi Prakash Srivastava, Shivaraman Pandey, Praveen Chandra Verma, Gauri Saxena

**Affiliations:** Department of Botany, University of Lucknow, Lucknow. INDIA; Plant Diversity, Systematics and Herbarium Division, CSIR-National Botanical Research Institute, Lucknow. INDIA; Plant Molecular Biology and Genetic Engineering Division, CSIR-National Botanical Research Institute, Lucknow. INDIA

**Author notes:** **Dr. Gauri Saxena**, Professor, Department of Botany, University of Lucknow, Lucknow-226007, Uttar Pradesh, INDIA.

**Keywords:** *Pinus*, Needles, Fascicles, Vascular bundle, Phylogeny

## Abstract

Pine is native to all continents and some oceanic islands of the northern hemisphere, chiefly in boreal, temperate or mountainous tropical regions; reaching its southernmost distribution below the equator in Southeast Asia. *Pinus* is divided into two subgenera, *Strobus*, and *Pinus* by the number of vascular bundles present in the needles. Comprehensive and detailed anatomy of needles in ten species of *Pinus* using nine anatomical traits was carried out. These morphological and anatomical traits supported the classification of the genus up to section level. It was observed that number of needles per fascicle varied along with other related traits such as thickness and width of vascular bundles, the diameter of resin ducts, the thickness of epidermis and thickness and width of endodermal cells that show remarkable variations among different species selected for the present study. The data can be used as a tool for identification and classification of *Pinus* upto genus and species level. We also found that similarity and differences in leaf anatomical traits supported the molecular phylogeny of *Pinus* conducted by several researchers.

## 1. Introduction

Order Pinales represents an outstanding group of gymnosperms and is omnipresent in terrestrial habitat. These monoecious woody plants usually grow naturally or have been introduced in both the hemispheres, mainly in the Northern hemisphere sometimes occurring in subtropical and tropical regions of Central America and Asia. They may form forest or co-exist with other trees (Farjon 1984 and 2005; Gaussen et al. 1993). They have received much attention in past few years because they form a major component of many temperate forests, and not only have ecological implications but are also of economical significance as a source of timber, pulp and paper, nuts, seeds, resins, construction materials, and other by-products. (Richardson and Rundel 1998).

*Pinus* is a popular tree plant in Indian Himalayas with high medicinal value and has played a significant role in maintaining health. *Pinus* is the largest genus in the family of coniferous trees with broad climate adaptability. *Pinus* has 110 species and is usually divided into two subgenera, *Strobus* (soft pines) and *Pinus* (hard pines), which are further divided into sections and subsections (Little and Critchfield 1969; Gernandt et al. 2005). The taxonomic history of *Pinus* was reviewed by Price et al.1998, that included morphology, anatomy, crossability, cytology, secondary metabolites, DNA and protein comparisons. The morphological traits that distinguish various species from one another in *Pinus* include characters like the length and width of needles, the number of needles per fascicle, arrangement and orientation of needles (pendulous or erect), and anatomical characteristics like epidermal cells, number and position of resin ducts, and number of vascular bundles in the needle (Gernandt et al. 2005).

It has been observed that traits of needles like shape, width, thickness, cuticle, thickness and width of epidermis, length and width of vascular bundles, resin canals and other morphological and anatomical characters of stem like wood, structure of axial tracheids, axial parenchyma change with environmental factors like temperature, light availability, and moisture content in the habitat where they grow since they are directly exposed to the environment ((Abrams and Kubiske, 1990., Dixit et. al. 2016). These morphological as well as anatomical differences could provide new information that can be used to establish phylogenetic relationship among various species. (Ghimire et. al. 2014). Although the needle structure of the common conifers like that of *Pinus* is comparatively well studied and known, a comprehensive treatment of the comparative histological organization of the needles is still lacking. The main focus of present work was to perform a detailed anatomical study of needles from native Indian and cultivated species of *Pinus* to make a detailed histological comparison of selected *Pinus* species. The data generated was further used to draw evolutionary relationship among these ten exotic and indigenous species of *Pinus* namely *P.merkusii, P.khasya, P.taeda, P.elliottii, P.echinata, P.thunbergii, P.patula, P.greggii, P.wallichiana*, and *P.roxburghii*.

## 2. Material and Method

### 2.1 Sample collection

Altogether 150 observation by analysing 30 plant samples (3 trees for each species and five needles per tree), comprising 10 pine species (Table 1) were collected using random block design (RBD) in September, 2016, from a cultivated population in the region of Ranikhet (located at 357 km NSE of New Delhi (Fig 1): latitude 29°39’52.2” (N); longitude 79°28’40.9” (E); altitude 1,727 m). The site is characterized by an average temperature of 14.4 °C, median rainfall (about 1575 mm of precipitation annually, and low soil fertility. A voucher specimen of all the species selected for study was deposited in the herbarium of the National Botanical Research Institute, Lucknow and identified.

**Table 1.** Selected species of *Pinus*.

| No. | Plant material | Sample code | Locality |
| --- | --- | --- | --- |
| 1. | *P.merkusii* * | **PM** | NW Himalayan Mtn. |
| 2. | *P.khasya* * | **PK** | NW Himalayan Mtn. |
| 3. | *P.taeda* | **PTd** | NW Himalayan Mtn. |
| 4. | *P.elliottii* | **PEI** | NW Himalayan Mtn. |
| 5. | *P.echinata* | **PE** | NW Himalayan Mtn. |
| 6. | *P.thunbergii* * | **PT** | NW Himalayan Mtn. |
| 7. | *P.patula* | **PP** | NW Himalayan Mtn. |
| 8. | *P.greggii* | **PG** | NW Himalayan Mtn. |
| 9. | *P.wallichiana* * | **PW** | NW Himalayan Mtn. |
| 10. | *P.roxburghii* * | **PR** | NW Himalayan Mtn. |

**\*Pine species native to India.**

**Figure 1.**
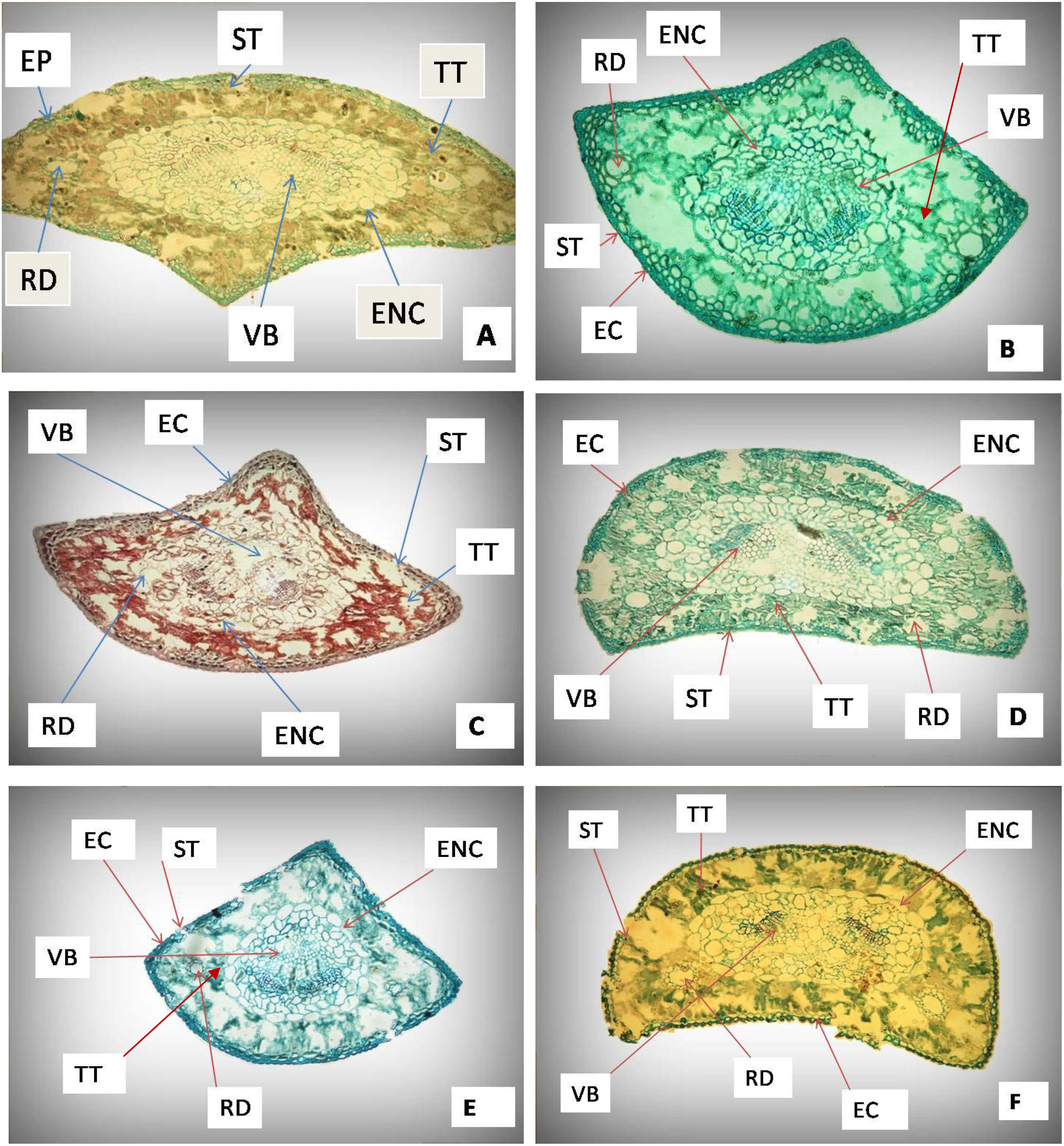
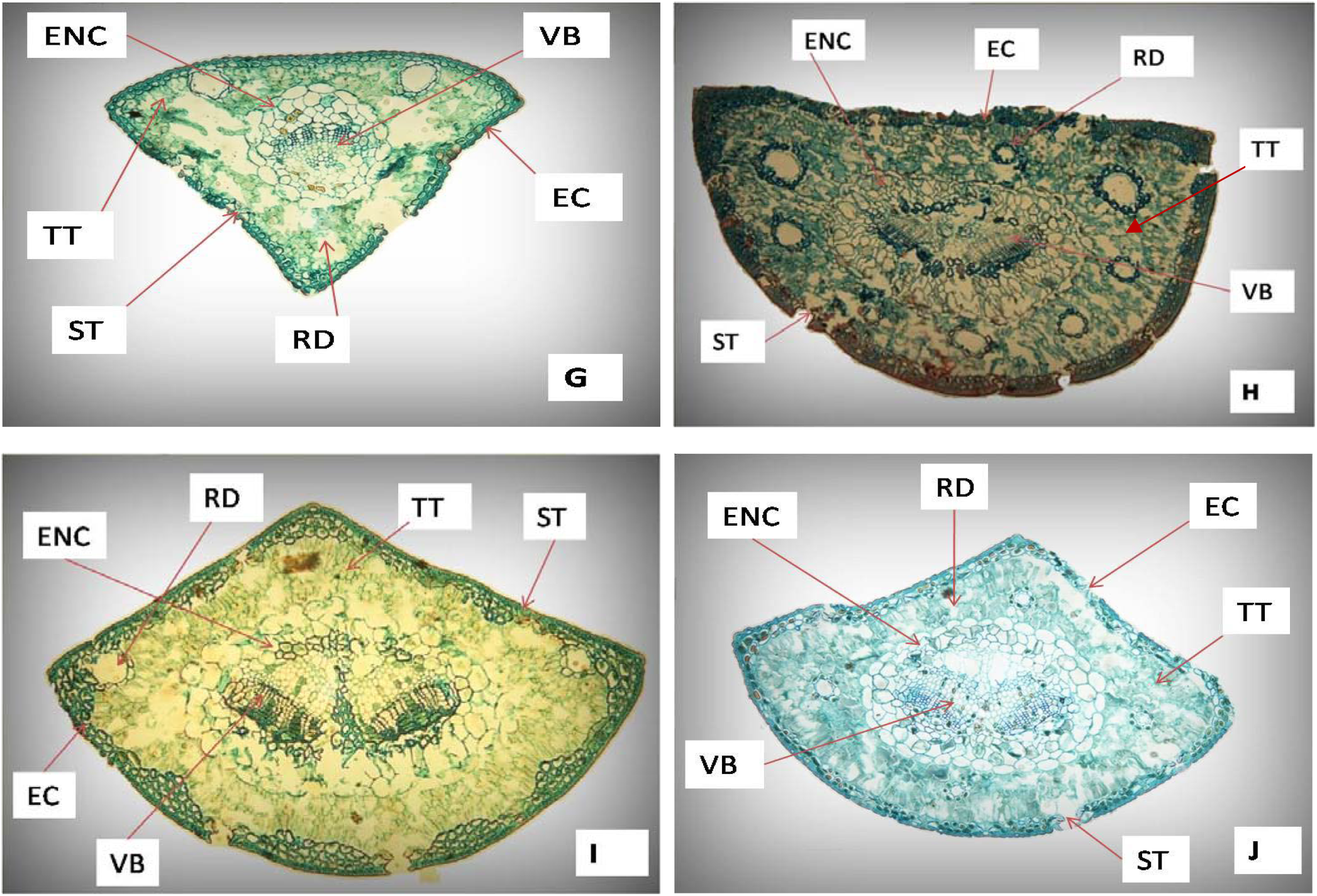
**A**;*P.merkusii*, **B**;*P.khasya*, **C**;*P.elliottii*, **D**;*P.echinata*, **E**;*P.taeda*, **F**;*P.patula*, **G**;*P.wallichiana*, **H**;*P.thunbergii*, I;*P.roxburghii*, **J**;*P.greggii*. [EC, Epidermal cell;ENC, Endodermal cell; RD,resin duct; TT,Transfusion tissue; ST, stomata; VB,vascular bundle]

### 2.2 Methodology for morpho-anatomical studies

Tree height was measured using stick method and other macroscopic and microscopic analysis was performed at the laboratory in the Department of Botany, University of Lucknow, Lucknow. Needle length was measured using a measuring scale, whereas other anatomical characteristics (Table 2; needle width, needle thickness, epidermis thickness, hypodermis thickness, and resin duct diameter) were observed (Fig. 1). under the microscope (Nikon Eclipse 80i). For morphological studies, parameters like a number of needles per fascicle and needle length were taken. The other morphological characters taken into account included the height of the plant and bark color. For anatomical studies, fresh plant needles were collected from *Pinus* species under study. The microtome sectioning (using Radical Di-cast Microtome, RMT-20A) and processing of needles was done according to the procedure given by Federica and Ruzin, 2000. Fresh needles were fixed in a formalin-acetic acid solution (50 ml 95% ethanol, 5 ml glacial acetic acid, 10 ml 37% formaldehyde and 35 ml distilled water) and kept for 24 hours for fixation. Before the final dehydration process, the fixed tissue was slightly warmed in 1% sodium hydroxide. In this step, needles were treated with an increasing gradient of a mixture of ethanol, tert-butanol, and water starting from 10% tert-butanol and ending at 100% tert-butanol. The samples were kept in each grade for a minimum of 35 minutes. After dehydration, the samples were transferred into wax containers where the wax was kept at 60-65 °C temperature in an oven. Molten wax was changed at a regular interval of 30 minutes with the addition of fresh wax. In this step, freshly molten wax was poured in the cubic space formed by attaching two L Blocks. Then the samples which were previously treated with wax were inserted vertically in the semisolid wax and wax block with the embedded tissue was allowed to cool followed by separation of L-Blocks. The wax cubes with the embedded specimen were then fixed to wooden blocks to get them attached to the microtome consequently. The attached wax blocks were then subjected to section cutting using microtome and sections were cut between 8μm-12μm depending upon the hardness of the samples.

**Table 2.** Needle anatomical traits analyzed.

| Abbreviation | Unit | Traits |
| --- | --- | --- |
| NT | µm | Needle thickness |
| NW | µm | Needle width |
| C+ET | µm | Cuticle and epidermal thickness |
| ECW | µm | Epidermal cell width |
| ENCT | µm | Endodermis cell thickness |
| ENCW | µm | Endodermis cell width |
| VBW | µm | Vascular bundle width |
| VBT | µm | Vascular bundle thickness |
| RCD | µm | Resin Canal Diameter |

### 2.3 Preparation of slides

The wax films of determined thickness were scooped out on a slide coated with a layer of a mixture of egg albumin and glycerol, which acts as an adhesive. The slide containing the paraffin film was then gently warmed and dipped in a jar containing xylene and was kept till all the wax gets solubilized in xylene. The slide was treated with alcohol solution starting from a concentration of 95%, then 90%, 80%, 70% and finally 50%. The slide was then stained with safranin solution (0.5%) and was kept for 12-15 minutes depending on nature and thickness of the sample. The slides were further treated with an increasing grade of ethanol starting from 50% and ultimately ending in 90% followed by staining with the fast green solution (0.1%). Slides were eventually washed with 95% ethanol to clear any excess stain in the section. They were dried, and the prepared sections were mounted on DPX and covered with a cover slip attentively avoiding any air bubble. Investigations and measurements of all selected anatomical traits were carried out on needles of all ten species of *Pinus* using the light microscope fitted with a Nikon digital camera in National Botanical Research Institute (NBRI). The slides of each of the studied plant parts were examined under a microscope; the eye piece lens was (×10) whereas the objective lenses were (×4 and ×20).

### 2.4 Statistical analysis

All samples were analyzed in triplicates, and their mean and standard deviation (SD) were calculated accordingly.Varaition in anatomical traits were compared by using one way analysis of variance (ANOVA), followed by Duncan multiple range test (DMRT) using SPSS16.0 software. The dendrogram was generated using the nearest neighbor method, squared Euclidean distance measure, based on differences between measurements of anatomical traits using Statgraphics Plus version 5.0 (Statistical Graphics Corporation, Princeton, NJ, USA). Principal components analysis (PCA) was applied to scale data and evaluate the underlying dimensionality of the variables and to elucidate the relationship among selected traits using Past v software.

## 3. Results

The results of the morphological (Table 3) and anatomical (Table 4 a & b; Fig. 1) studies on various traits of *Pinus* needles collected from a wild population in the region of Northwest Himalayas in the Indian state of Uttarakhand have been summarized below.

**Table 3.** Morphological trait analysis in selected species of *Pinus*

| *Pinus* species | Height<br>(in feet) | Bark color | No of needle<br>per fascicles | Size of needle<br>(in Inches) |
| --- | --- | --- | --- | --- |
| *Pinus thunbergii* | 90-130 | Greyish brown | 2 | 4.2±1.3 |
| *Pinus roxburghii* | 150-180 | Dark grey | 3 | 11.8±2.7 |
| *Pinus wallichiana* | 50-150 | Greyish brown | 5 | 6.1±1.9 |
| *Pinus merkusii* | 45-60 | Grey to brown | 2 | 8.5±1.6 |
| *Pinus khasya* | 100-150 | Light brown | 3 | 7.3±1.2 |
| *Pinus taeda* | 90-110 | Reddish brown | 3 | 5.9±1.1 |
| *Pinus echinata* | 80-120 | Reddish | 2 | 4.0±1.0 |
| *Pinus gregii* | 40-50 | Grey | 3 | 5.2±1.7 |
| *Pinus patula* | 40-60 | Reddish brown | 3 | 7.9±1.7 |
| *Pinus elliottii* | 60-100 | Greyish brown | 2 | 8.3±1.3 |

**Table 4(a).**
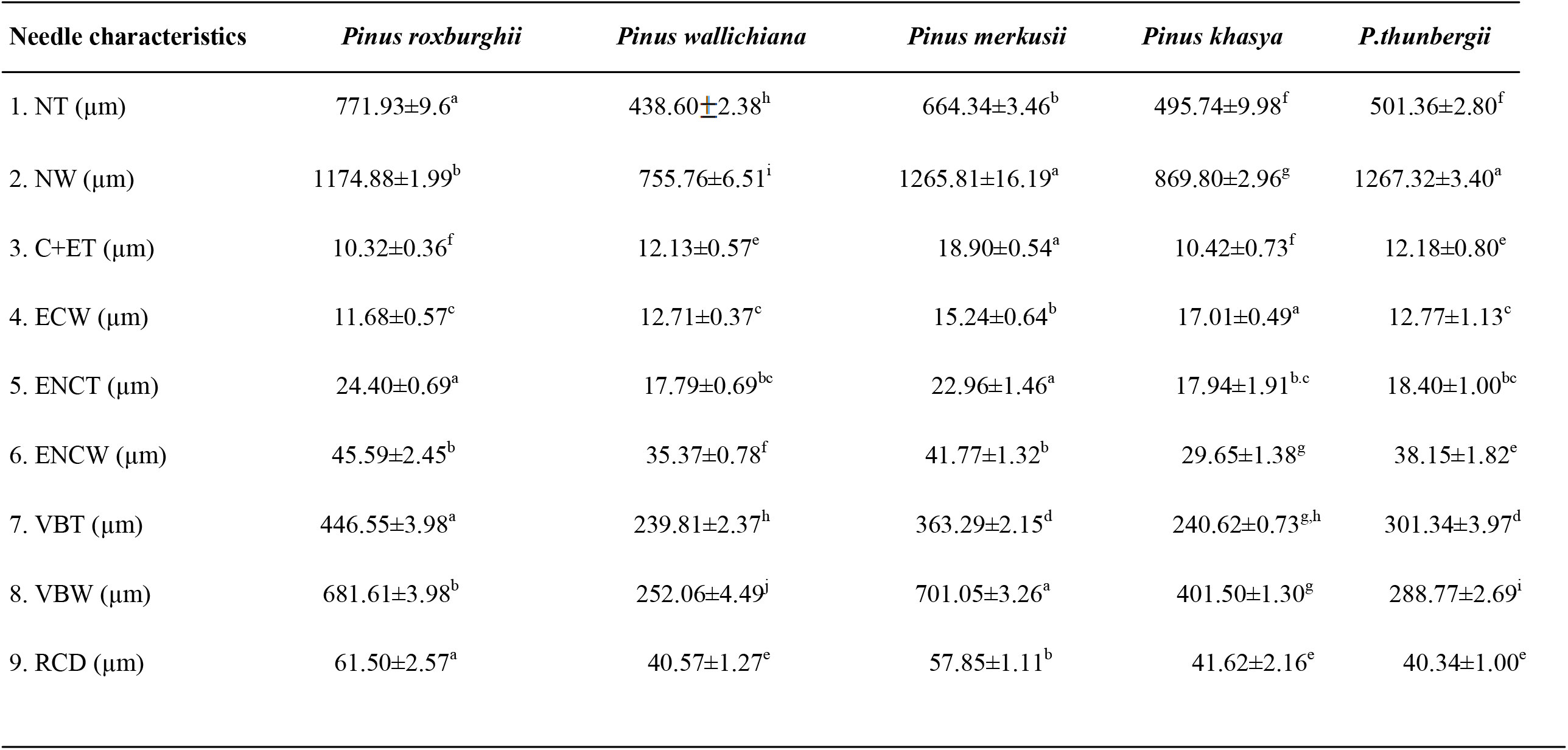
Anatomical traits of needles in selected *Pinus* species. Mean ± standard deviation (SD) (n = 3). Letters indicate significant differences between different traits each parameter separately using Duncan multiple range tests (DMRTs) at significant p b .05 level (ANOVA analysis, n = 3).NT, needle thickness; NW, needle width: C+ET, cuticular + epidermal thickness; ECW, epidermis cell width; ENCT, endodermis cell thickness; ENCW, endodermis cell width; VBT, vascular bundle thickness; VBW, vascular bundle width; RCD, resin canal diameter.

**Table 4(b).** Anatomical traits of needles in selected *Pinus* species. Mean ± standard deviation (SD) (n = 3). Letters indicate significant differences between different traits each parameter separately using Duncan multiple range tests (DMRTs) at significant p b .05 level (ANOVA analysis, n = 3).NT, needle thickness; NW, needle width: C+ET, cuticular + epidermal thickness; ECW, epidermis cell width; ENCT, endodermis cell thickness; ENCW, endodermis cell width; VBT, vascular bundle thickness; VBW, vascular bundle width; RCD, resin canal diameter.

| Needle characteristics | *Pinus taeda* | *Pinus echinata* | *Pinus greggii* | *Pinus patula* | *P.elliottii* |
| --- | --- | --- | --- | --- | --- |
| 1. NT ($\mu\text{m}$) | 645.66±2.74<sup>c</sup> | 578.44±2.59<sup>e</sup> | 607.09±4.61<sup>d</sup> | 485.27±4.10<sup>g</sup> | 502.48±3.05<sup>f</sup> |
| 2. NW ($\mu\text{m}$) | 1024.17±5.41<sup>c</sup> | 972.70±2.30<sup>e</sup> | 847.96±3.44<sup>h</sup> | 899.49±2.40<sup>f</sup> | 1011.89±2.57<sup>d</sup> |
| 3. C+ET ($\mu\text{m}$) | 13.84±0.25<sup>d</sup> | 17.37±0.33<sup>b</sup> | 19.42±0.43<sup>a</sup> | 14.96 ±0.09<sup>c</sup> | 11.68±0.33<sup>e</sup> |
| 4. ECW ($\mu\text{m}$) | 12.40±0.64<sup>c</sup> | 14.83±0.85<sup>b</sup> | 11.68±0.62<sup>c</sup> | 16.80±0.30<sup>a</sup> | 15.08±0.51<sup>b</sup> |
| 5. ENCT ($\mu\text{m}$) | 16.74±0.83<sup>c</sup> | 23.32±0.98<sup>a</sup> | 19.23±0.49<sup>b</sup> | 11.72±0.46<sup>e</sup> | 14.00±0.27<sup>d</sup> |
| 6. ENCW ($\mu\text{m}$) | 51.61±0.78<sup>a</sup> | 43.04±1.16<sup>cd</sup> | 41.00±1.09<sup>d</sup> | 46.65±0.79<sup>b</sup> | 43.39±1.16<sup>cd</sup> |
| 7. VBT ($\mu\text{m}$) | 324.69±3.16<sup>b</sup> | 269.88±1.16<sup>e</sup> | 266.11±4.96<sup>e</sup> | 255.71±2.87<sup>f</sup> | 245.69±1.26<sup>g</sup> |
| 8. VBW ($\mu\text{m}$) | 481.63±1.91<sup>f</sup> | 603.12±2.70<sup>c</sup> | 390.16±1.34<sup>h</sup> | 580.29±0.94<sup>c</sup> | 595.22±2.55<sup>d</sup> |
| 9. RCD ($\mu\text{m}$) | 55.07±1.15<sup>bc</sup> | 55.31±0.53<sup>c</sup> | 50.21±1.48<sup>d</sup> | 53.84±0.74<sup>c</sup> | 34.18±1.12<sup>f</sup> |

### 3.1 Morphological traits analysis

The analyzed populations of species were significantly different with respect to most of the selected traits. Maximum difference was observed in the height of plants which varied from an average of approximately 50 ft. In *Pinus merkusii* to an average of 155 ft. in *Pinus roxburghii*. Similarly, a number of needles per fascicles varied from about 2 to 5 most common being 3 and length of needles varied from 3 inches as in *Pinus echinata* and *Pinus thunbergii* to a maximum of about 13 inches in *Pinus roxburghii*. Variations were also observed in bark colour of identified *Pinus* species which varied from light brown in *Pinus khasya* to dark grey in *Pinus roxburghii*.

### 3.2 Anatomical traits analysis

A total of nine anatomical traits were taken into consideration, including needle thickness (NT); needle width (NW); cuticle and epidermal thickness (C+ET); epidermal cell width (ECW); endodermis cell thickness (ENCT); endodermis cell width (ENCW); vascular bundle thickness (VBT); vascular bundle width (VBW); resin canal diameter (RCD). Results of anatomical studies (Table 4 a & b) in needles of selected species of *Pinus* suggested significant variations in terms of needle width, needle thickness, vascular bundle width and thickness, thickness and width of epidermal cell and diameter of resin duct. Needle thickness was maximum in *P. roxburghii* while minimum in *P. wallichiana*. Among exotic species, needle thickness was maximum in *P.taeda*. The width of the needle was maximum in *P.thunbergii* while minimum in *P.wallichiana*. Among native species, maximum needle width was reported in *P.armandi*. Similarly, C+ET thickness was maximum in *P.greggii* while it was least in *P.roxburghii*. Among native species, it was maximum in *P.armandi*. As far as the width of epidermal cells was concerned, it was maximum in *P.khasya* while minimum in *P.greggii*. Among native species, maximum width was reported in *P.patula*. The thickness of endodermal cells was maximum in *P.roxburghii* while minimum in *P.patula*. Among exotic species, it was maximum in *P.echinata*. The width of endodermal cells was maximum in *P.taeda* while minimum in *P.khasya*. Among native species, the maximum width of endodermal cells was observed in *P.roxburghii*. Two important vascular bundle traits were taken into considerations viz. vascular bundle thickness and vascular bundle width. Among native species, former was reported maximum in *P.roxburghii* while minimum in *P.wallichiana* and later was reported maximum in *P.merkusii* while minimum in *P.wallichiana*. Among exotic species, *P. taeda* showed maximum value for vascular bundle thickness, and it was minimum for *P.elliottii* while vascular bundle width was maximum in *P.echinata* and it was minimum for P.greggii. The diameter of resin duct was maximum in *P.roxburghii* while minimum in *P.elliottii*. Among exotic species, *P.echinata* showed the maximum diameter of resin duct.

Cluster analysis was conducted using all the anatomical traits understudy for selected species of pine needles (Fig. 2). It displayed similarities between *P.merkusii* and *P.roxburghii* and between *P.wallichiana* and *P.khasya*. The results of the Principal component analysis (PCA) in the selected pine needles were obtained from nine anatomical characteristics (Fig. 3); 90 observations for each trait were processed in the correlation matrix. Each observation represented the average value of the properties analyzed in three needles per tree. PCA showed that the first two axes represent 65.76% of the information.

**Fig.2.**
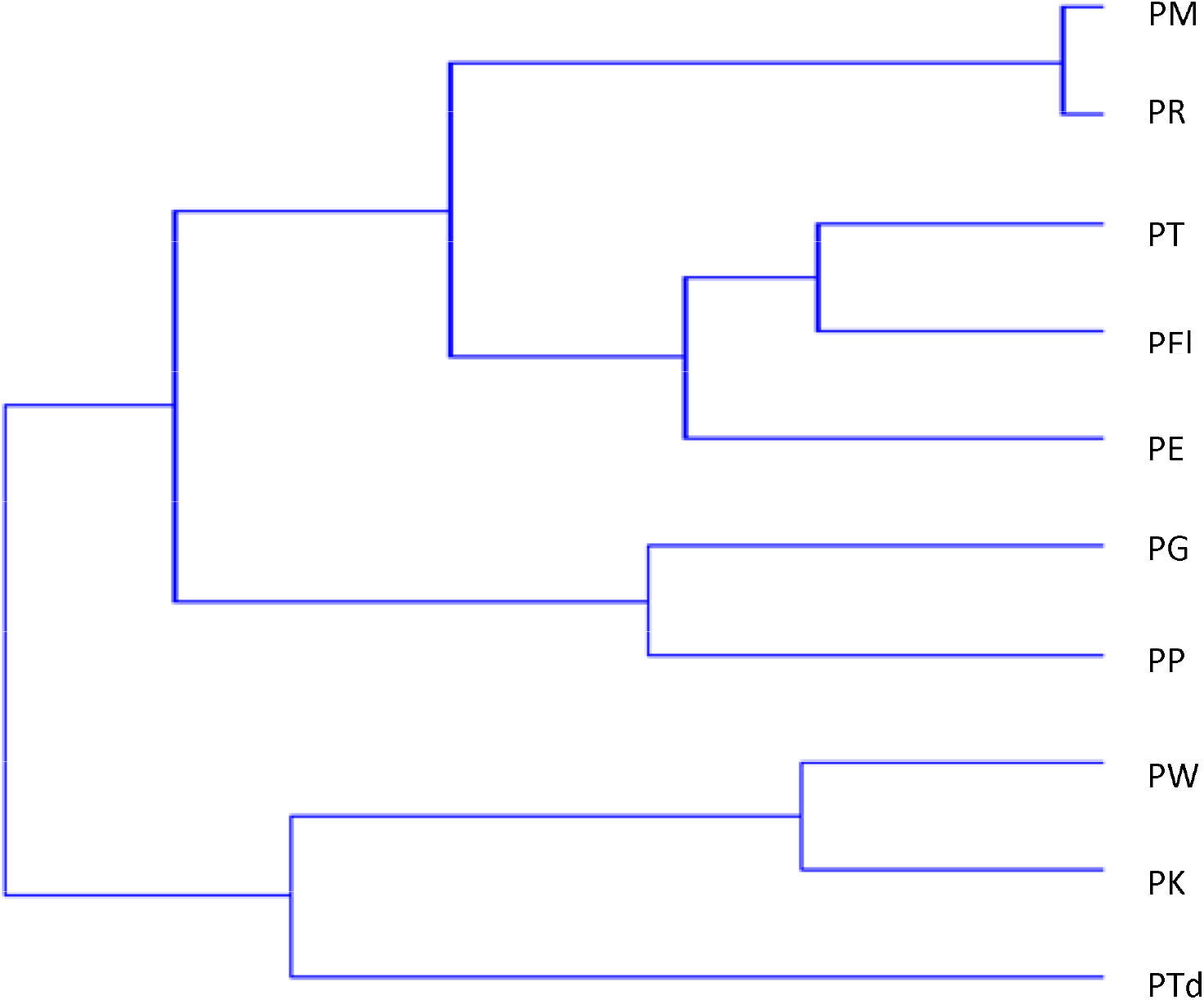
Dendrogram of nine morpho-anatomical properties of selected species of *Pinus* based on a “nearest neighbor method” (squared Euclidean distance).

**Fig 3.**
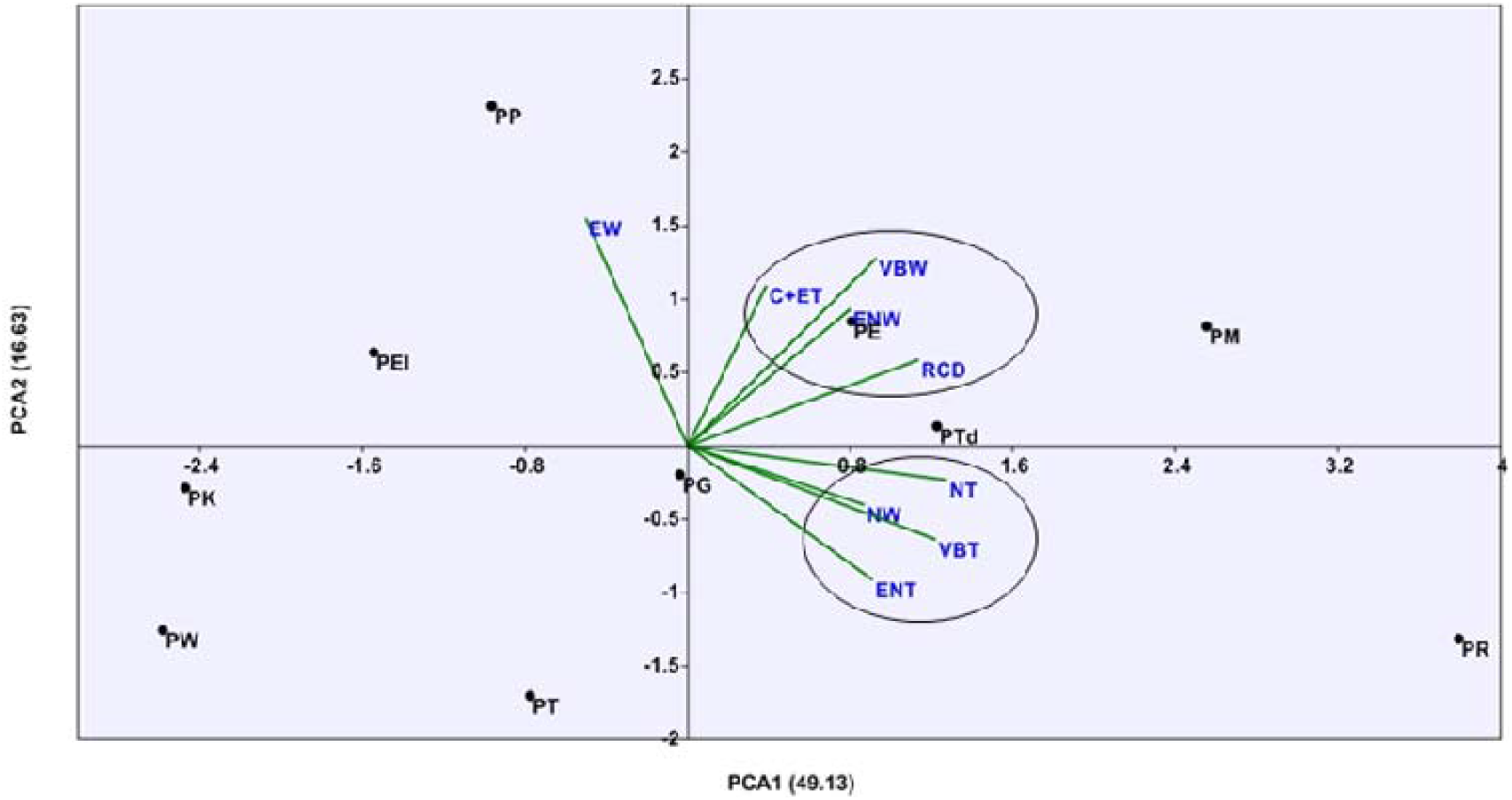
PCA of nine morpho-anatomical properties of selected species of *Pinus*.

**Fig. 4.**
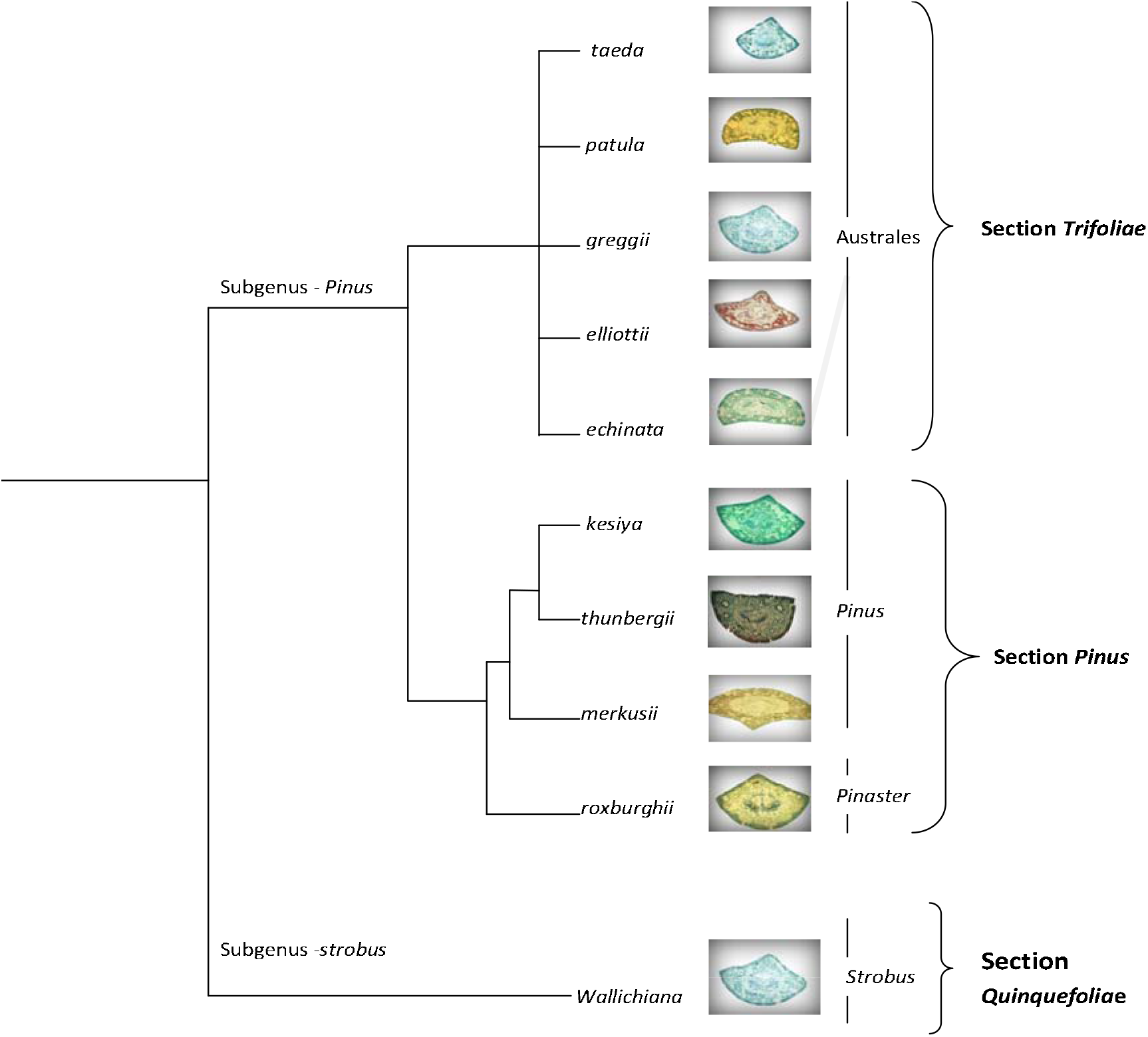
Phylogenetic tree showing the needle structure of selected Pinus species (modified from Gernandt et al. 2005)

## 4. Discussion

Pines differ from other members of family Pinaceae and easily characterized by their dimorphic shoots that include long as well as short shoots called fascicles. These fascicles bear long, narrow needle-like leaves mostly present in groups of two to five. The species in the three sections and three subsections included in this study had two, three, or five needles per fascicle (Table 2). Two species from section *Trifoliae (P. taeda and P. echinata)* had two needles per fascicle, whereas three taxa from section *Trifoliae* (*P.greggii, P.patula*, and *P.elliottii*) had three needles per fascicle. On the other hand, the two species of section *Pinus (P.thunbergii* and *P.merkusii)* had two needles per fascicle, whereas two taxa from section Pinus *(P.roxburghii* and *P.khasya)* had three needles per fascicle. Only taxa from section *Quinquefoliae*, i.e. *P.wallichiana* had five needles per fascicles. Within section *Trifoliae* all five selected species belong to subsection *Australes* while in section *Pinus* out of four selected species three belong to subsection *Pinus*, and one belongs to subsection *Pinaster*. Only species from section *Quinquefoliae* belong to subsection *Strobus*. From the above discussion, we can conclude that the number of needles per fascicle is a useful tool for recognizing species up to subsection level. The number of needles per fascicle has evolutionary significance too, as five needles per fascicle are considered to be a primitive character within *Pinus* as compared to two or three needles per fascicle (Kaundun and Lebreton 2010).

Out of the ten taxa examined in our study, nine belonged to subgenus *Pinus* and one to subgenus Strobus. In terms of the internal anatomical structure of the needle, two subgenera were easily distinguishable by the number of vascular bundles, as species of subgenus *Pinus* have two fibrovascular bundles per needle and those of subgenus Strobus have only single fibrovascular bundles. Further presence of two versus one vascular bundle within a single bundle sheath has proven to be an important diagnostic feature for differentiating subgenera *Strobus* from subgenera *Pinus* within genus Pinus (Gernandt et al. 2005; Eckenwalder 2009).

The morpho-anatomical parameters across the populations also form an important attribute to assess growth performance and biomass (Jugrana et al., 2013). In *Pinus*, needles are one of the most vigorous assimilatory organs, having important effects on plant physiology as well as ecological adaptability. Although most morphological and anatomical traits of needles remain stable at the species level, researchers have demonstrated that genetic variations do exist within them in general. Further, researchers have also discovered the adaptive features of needle traits in the environment. (Nobis et al. 2012; Legoshchina et al. 2013; Xing et al. 2014). In our study, we found a high level of morphological variability among native and exotic species as well as within their population particularly needle traits like needle length that shows a higher degree of variation. Same is also true for plant height and bark colour. Further, the individual has a specific norm of reactions to environmental factors and has a capacity for certain morphological modifications within a specific range.

In our study, we have selected nine anatomical traits including two traits from vascular system, i.e. vascular bundle width and vascular bundle thickness to see implications of these traits on the phylogeny of genus *Pinus*, and we encountered significant variations among populations that indicated large genetic differences (Table 4 a & b). However growing scientific evidence have shown that change in the internal structures of needles may be an outcome of climate change (Mao and Wang 2011). Both, correlation studies and PCA analysis based on observation of such traits showed relationship within vascular bundle traits, epidermal traits, and cross-section area traits that significantly varied and demonstrated that each part of the needle was relatively independent. It is interesting to observe that all exotic species (except *P.taeda*) show similarities among each other and exhibit variations when compared with native species (Fig. 2). Comparison between our dendograms based on a “nearest neighbor method” (squared Euclidean distance) using nine anatomical traits and that given by Gerandt et. al.,2005 (Fig.4) using chloroplast DNA for molecular phylogeny confirms the similarity between *P.greggii* and *P.patula* as well as between *P.elliottii* and *P.echinata*. It also confirms the similarity between *P.merkusii, P.roxburghii* and *P.thunbergii*. However position of *P.wallichiana, P.khasya* and *P.taeda* shows variability when their relationship with other species was studied using anatomical traits, to that of molecular phylogeny (Fig. 2 & 4). Our results are comparable to other researches on molecular phylogeny where chloroplast based markers have been used (Wang et al.,1999; Leon et. al., 2013; Olsson et. al.,2018).PCA visualizes that NW (Needle width), NT (Needle thickness), VBT (Vascular bundle thickness) and ENT (Endodermal thickness) show correlation in *P.roxburghii* and *P.taeda* while C+ET (Cuticular plus epidermal thickness), VBW (Vascular bundle width), ENW (Endodermal width) and RCD (Resin canal diameter) show correlation in *P.merkusii, P.echinata, P.taeda*.

## CONCLUSION

We found that variable needle anatomical traits exhibit great adherence to the molecular phylogeny of *Pinus* also attempted through chloroplast gene sequences and other markers earlier and provided reasonable evidence for classifying the genus upto subgenera, sections, and subsections level. However a large number of *Pinus* species are still anatomically not well studied or lack detailed anatomical explanations. The micro-measurement of various anatomical traits and other parameters like number and position of resin ducts, the position of vascular bundles, shape, and structure of leaves in cross section have great systematic value and are important for phylogenetic studies and classification of the genus *Pinus*. Further studies involving as many species as possible, including all subgenus, sections and subsections, are highly recommended for establishing a database for a full proof classification and identification of this genus.

## ACKNOWLEDGMENT

We are grateful to the University Grant Commission (UGC, India) for financial support provided by a grant to Lav Singh (Ref: 201314-NETJRF-10212-83). The author is also thankful to Head, Department of Botany, Lucknow University for providing necessary research facilities

## CONFLICT OF INTEREST

The authors declare no conflict of interest.

## References

Abrams, M. D. and M. E. Kubiske., 1990. Leaf structural characteristics of 31 hardwood and conifer tree species in central Wisconsin: Influence of light regime and shade-tolerance rank. Forest Ecology and Management 31: 245–253.

Dixit, P., Singh, L., Verma, P.C., and Saxena, G., 2016. Altitudinal Influences on Leaf and Wood Anatomy and its Ecological Implications in Cephalotaxus griffithii of Indian Himalayas. J. Biol. Chem. Research. Volume 33 (1) 2016 Pages No. 388–399

Jugrana, A.K., Bhatt, I.D., Rawal, R.S., Nandi, S.K., Pande, V., 2013. Patterns of morphological and genetic diversity of *Valeriana jatamansi* Jones in different habitats and altitudinal range of West Himalaya, India. Flora. 208, 13–21. https://doi.org/10.1016/j.flora.2012.12.003

Eckenwalder, J.E., 2009. Conifers of the World. Timber Press, Portland

Farjon, A., 1984. Pines: drawings and descriptions of the genus Pinus. EJ Brill and W Backhuys, Leiden

Farjon, A., 2005. Pines: drawings and description of the genus Pinus, 3rd edn. Brill, Leiden.

Federica, B., Ruzin, S.E., 2000. Plant Micro technique and Microscopy. Oxford University Press, New York.

Gaussen, H., Heywood, V.H., Chater, A.O., 1993 Pinus L. In: Tutin, T.G., Burges, N.A., Chater, A.O., Edmondson, J.R., Heywood, V.H., Moore, D.M., Valentine, D.H., Walters, S.M., Webb, D.A.,(eds) Flora Europaea, vol 1, 2nd edn. Cambridge University Press, Cambridge, 40–44.

Gernandt, D.S., Lopez, G.G., Garcia, S.O., Liston, A., 2005. Phylogeny and classification of *Pinus*. Taxon 54, 29–42.

Ghimire, B., Kim, M., Lee, J.H., Heo, K., 2014. Leaf anatomy of Pinus thunbergii Parl. (Pinaceae) collected from different regions of Korea. Korean Journal of Plant Taxonomy 44 (2), 91–99.

Kaundun, S.S., Lebreton, P., 2010. Taxonomy and systematics of the genus *Pinus* based on morphological, biogeographical and biochemical characters. Plant Syst Evol 284, 1–15

Legoshchina, O., Neverova, O., Bykov, A., 2013. Variability of the anatomical structure of *Picea* obovata Ledeb. Needles under the influence of emissions from the industrial zone of Kemerovo. Contemp Probl Ecol 6, 555–560

Leon, S.H., Gernandt, D.S., Rosa, J.A., Barbolla, L.Z., 2013. Phylogenetic Relationships and Species Delimitation in *Pinus* Section *Trifoliae* Inferrred from Plastid DNA. PLOS ONE (8)

Little, E.L., Critchfield, W.B., 1969. Subdivision of the genus *Pinus* (Pines). USDA Forest Service Miscellaneous Publication 1144, Washington, DC

Mao, J.F., Wang, X.R., 2011. Distinct niche divergence characterizes the homoploid hybrid speciation of *Pinus densata* on the Tibetan Plateau. Am Nat 177, 424–439.

Olsson, S., Grivet, D., Vian, J.C., 2018. Species-diagnostic markers in the genus *Pinus*: evaluation of the chloroplast regions matK and ycf1. Forest system 27,1–11.

Price, R.A., Liston, A., Strauss, S.H., 1998. Phylogeny and systematics of Pinus. In: Richardson DM (ed) Ecology and biogeography of Pinus. Cambridge University Press, Cambridge, 49–68.

Nobis, M.P., Traiser, C., Roth, N.A., 2012. Latitudinal variation in morphological traits of the genus *Pinus* and its relation to environmental and phylogenetic signals. Plant Ecol Divers 5,1–11

Richardson, D.M., Rundel, P.W., 1998. Ecology and biogeography of *Pinus*: an introduction. In: Richardson DM (ed) Ecology and biogeography of Pinus. Cambridge University Press, Cambridge.

Wang, X.R., Tsumura, Y., Yoshumaru, H., Nagasaka, K., Szmidt, A.E., 1999. Phylogenetic relationships of Eurasian Pines (*Pinus*, Pinaceae) based on Chloroplast Rbc L, MATK, Rpl20-Rps18 SPACER, and TRNV intron sequences. American Journal of Botany. 86, 1742–1753.

Xing, F.Q., Mao, J.F., Meng, J.X., Dai, J.F., Zhao, W., Liu, H., Xing, Z., Zhang, H., Wang, X., Li, Y., 2014. Needle morphological evidence of the homoploid hybrid origin of *Pinus densata* based on analysis of artificial hybrids and the putative parents, *Pinus tabuliformis* and *Pinus yunnanensis*. Ecol Evol 4,1890–1902.

